# Benchmarking HelixFold3-Predicted Holo Structures for Relative Free Energy Perturbation Calculations

**DOI:** 10.1101/2024.10.27.620454

**Authors:** Kairi Furui, Masahito Ohue

## Abstract

AlphaFold2 demonstrated remarkable capabilities for protein structure prediction. However, it is limited to downstream tasks, such as ligand docking and free energy calculations, as it cannot predict holo structures with bound ligands. AlphaFold3, a state-of-the-art protein structure prediction model, can predict the binding structures of complexes with proteins, nucleic acids, small molecules, ions, and modified residues with cutting-edge performance. However, AlphaFold3 does not currently provide access to some functions, such as the prediction of protein-ligand complex structures. To reduce the enormous costs in early small molecule drug discovery, verifying the utility of protein-ligand complex prediction methods, such as AlphaFold3, in downstream tasks like free energy perturbation calculations is crucial. In this study, we evaluated HelixFold3, designed to emulate AlphaFold3, in predicting holo and apo structures’ complex formations and examined its utility in free energy perturbation calculations.

Regarding the complex structure prediction performance of the 8 targets from Wang *et al*.’s FEP benchmark, HelixFold3 showed superior performance to AlphaFold2 and existing methods. Predicting a holo structure rather than an apo structure resulted in higher binding site prediction accuracy. Furthermore, using HelixFold3 predicted structures in practical situations, where binding free energies of all derivatives were estimated, both structures achieved accuracies comparable to crystal structures. Additionally, novel derivatives not included in the training data were accurately predicted, demonstrating that free energy calculations using these novel structures are sufficiently usable.

## Introduction

Protein structure prediction is vital, and DeepMind’s AlphaFold series^1–3^ is a prime example of successful deep learning applications in science. The latest AlphaFold3^3^ demonstrates state-of-the-art prediction for the binding structures of complexes with small molecule ligands, nucleic acids, ions, and modified residues. However, as of October 2024, AlphaFold3’s implementation has not been released, with only a limited online server (https://alphafoldserver.com/) available for structure prediction. Liu *et al*. developed HelixFold3,^4^ a model with comparable performance to AlphaFold3, and released its code and models. HelixFold3 provides capabilities restricted in AlphaFold3, such as protein-ligand complex structure prediction, and is expected to perform similarly in these tasks.

Since the emergence of AlphaFold2, studies have evaluated not only protein structure prediction performance but also the quality of predicted structures in downstream tasks, such as free energy perturbation (FEP) calculations and virtual screening.^5–8^ Beuming *et al*. performed FEP calculations using FEP+^5^ with structures predicted by restricting structural templates and sequence information to 30% identity, especially when structural information is limited, and evaluated the accuracy.^9^ Their results indicated that the performance of ΔΔ*G* calculations using AlphaFold2 structures was comparable to that using crystal structures.

However, they only performed calculations for a carefully selected subset of perturbations due to computational resource limitations. Therefore, the performance in real scenarios of calculating the entire dataset was not evaluated. Additionally, the predicted apo structures were refined by protein-ligand complex refinement and manual adjustment to resolve clashes, suggesting that using AlphaFold2’s predicted apo structures directly for FEP calculations was inappropriate. Moreover, Díaz-Rovira *et al*.^7^ and Holcomb *et al*.^8^ reported that virtual screening using AlphaFold2 predicted structures resulted in lower screening performance compared to using crystal structures. Therefore, although AlphaFold2 shows high prediction performance for proteins overall, it is challenging to accurately predict more detailed structures, such as binding sites, suggesting limitations in its predicted structures for downstream tasks.

Due to AlphaFold2’s limitations, accurate prediction of binding sites with AlphaFold3’s capabilities is crucial for practical drug discovery campaigns. The ligand docking performance of AlphaFold3 has been evaluated using the PoseBusters^10^ benchmark. Based on the percentage of protein-ligand pairs with a ligand root mean square deviation (RMSD) of less than 2 Å, it achieved a higher success rate than existing docking methods such as AutoDock Vina^11^ and RoseTTAFold All-Atom.^12^ However, the accuracy of binding site predictions and the utility of the predicted structures in downstream tasks, such as free energy calculations and virtual screening, have not been thoroughly evaluated. If the predicted complex structures can not only predict the ligand’s binding position and pose but also reflect the actual holo structure’s binding pocket, they can be widely utilized in downstream tasks more than AlphaFold2 did.

In this study, we evaluated the usefulness of the predicted structures for FEP calculations to verify HelixFold3’s protein-ligand complex prediction performance in practical downstream tasks. Using Wang *et al*.’s FEP benchmark dataset, ^5^ we assessed the prediction accuracy of binding free energies using Flare FEP^13,14^ with HelixFold3’s predicted holo and apo structures. Unlike Beuming *et al*.,^9^ we performed FEP calculations on the entire dataset, not just a subset, to evaluate realistic scenario performance. It is also essential to determine whether HelixFold3 can predict protein-ligand complex structures with unknown ligands not present in the PDB dataset, and if these structures are useful. Thus, we assessed HelixFold3’s generalizability and robustness by predicting the complexes of all derivatives in Wang *et al*.’s benchmark set. For each target, we selected one predicted complex structure with its derivative and evaluated its FEP performance.

## Materials and Methods

### Evaluation of FEP Performance

To evaluate FEP performance using HelixFold3 predicted structures, we used 8 targets from Wang *et al*.’s FEP benchmark dataset. ^5^ Using 3 structures: the crystal structure, HelixFold3’s predicted holo structure, and the predicted apo structure, we constructed perturbation maps using Cresset Flare FEP V8.0.2^13,14^ and performed FEP calculations. Default perturbation maps were constructed using Flare FEP’s perturbation map construction function. We introduced intermediates to facilitate complex transformation links in the perturbation map, reconstructing the map when the link score, ^15^ indicating compound transformation complexity, was 0.6 or less. Table 1 details each target’s crystal structure and the constructed perturbation map’s statistics (number of ligands, introduced intermediates, and perturbation links). The same perturbation map was used for each target protein. All structures predicted by HelixFold3 were superimposed on the crystal structure. The ligand pose used was not the predicted pose from HelixFold3 but the crystal ligand pose, and the derivative poses were aligned to the crystal structure using Cresset Flare’s ligand alignment function.^14^

**Table 1:** Details of Wang *et al*.’s FEP benchmark and statistics of the constructed perturbation maps.

| Target | PDB | Chains | Crystal ligand | #Ligands(#Intermediates) | #Links |
| --- | --- | --- | --- | --- | --- |
| BACE | 4DJW | A | 0KP | 36(3) | 57 |
| CDK2 | 1H1Q | A | 2A6 | 16(2) | 23 |
| JNK1 | 2GMX | A | 877 | 21(0) | 27 |
| MCL1 | 4HW3 | A | 19G | 42(1) | 63 |
| P38 | 3FLY | A | FLY | 34(8) | 64 |
| PTP1B | 2QBS | A | 024 | 23(2) | 35 |
| Thrombin | 2ZFF | A | 53U | 11(0) | 14 |
| TYK2 | 4GIH | H | 0X5 | 16(1) | 23 |

### FEP Protocol

The FEP calculations were performed using the default settings of Cresset Flare FEP V8.0.2.^13,14^ The AMBER FF14SB force field^16^ modeled the proteins. The General AMBER Force Field 2 (GAFF2)^17^ was applied to the ligand force field, and AM1-BCC^18^ was adopted for partial charges. Further, the TIP3P^19^ explicit water solvent model was used.

Energy minimization was performed to remove bad contacts, with a tolerance of 0.25 kcal/mol for the free system and 2.50 kcal/mol for the bound system, using up to 1,000 iterations. Following minimization, the systems were gradually heated under NVT conditions to reach the target temperature of 298 K. Each simulation step was conducted with a 4.0 fs time step, and the system was equilibrated for 500 ps, followed by a 4 ns production run.

**Stage 1:** Strong positional restraints (strength of 2.00) were applied to both protein and ligand heavy atoms to maintain structural integrity during initial equilibration.

**Stage 2:** Restraints were reduced to the protein backbone atoms and ligand heavy atoms, allowing side chains to relax.

**Stage 3:** Very weak restraints (strength of 1.00) were applied to the protein backbone atoms, with the ligand unrestrained, to allow further relaxation of the system.

The total equilibration time was approximately 500 ps, with each stage lasting 100–200 ps.

This multistage equilibration protocol ensured that the system reached equilibrium before production runs.

Production runs were conducted in the NPT ensemble using a 4.0 fs time step. Bonds involving hydrogen atoms were constrained, except for those in the perturbed regions during alchemical transformations.

In Flare FEP, the number of *λ* windows (denoted as *λs*) for each alchemical transformation is automatically determined at runtime based on convergence criteria, with the minimum number of *λs* set as *minλ*.

### HelixFold3

We compared inference results using HelixFold3 with the protein and binding ligand as inputs and with only the protein as input. Input data were obtained from the PDB ID, chains, and crystal ligands listed in Table 1. HelixFold3 model parameters are publicly available. During inference, we used the full Big Fantastic Database (BFD) sequence database ^1,20,21^ instead of HelixFold3’s default reduced BFD database.

Structures up to February 2024 from the Protein Data Bank^22^ were used as templates for HelixFold3 prediction. To evaluate model quality, 5 inferences were made for each target to confirm the variety of conformations.

### Evaluation of Predicted Structures in Derivative Sets

The 8 targets of Wang *et al*.’s FEP benchmark already have crystal structures in the PDB and may be included in HelixFold3’s training data. Thus, even if complex structures can be predicted for these targets, generalization performance cannot be guaranteed. We evaluated the accuracy of predicting the holo structure for each derivative of Wang *et al*.’s FEP benchmark. The derivative set included many ligands not registered in the PDB, allowing evaluation of HelixFold3’s ability to predict complex structures with unknown ligands. Then, we selected one complex from the predicted complexes with a different binding pocket structure than the predicted complex with the crystal ligand and evaluated the performance of the free energy calculation. This study aimed to evaluate whether HelixFold3’s predicted holo structures are useful for FEP calculations in a blind situation where true complex structures are not trained.

For each target, complex structures were predicted by HelixFold3 for the number of ligands listed in Table 1. Only the structure with the highest HelixFold3 ranking score was selected for each derivative. The derivative ligand complex structure is not necessarily similar to the complex structure of the crystal ligand; however, it is assumed that no significant structural changes occur in the crystal ligand’s complex structure.

### Comparison of Structure Prediction Methods

Among the existing structure prediction methods, ColabFold,^23^ RoseTTAFold All-Atom (RoseTTAFold-AA),^12^ and Umol^24^ were used to compare HelixFold3’s complex structure prediction performance. ColabFold combines AlphaFold2 with MMseqs2’s^25^ fast homology search. In this study, ColabFold 1.5.5 (compatible with AlphaFold 2.3.2^2^) used all default parameters and performed inference of 5 AlphaFold2 models. Umol predicts protein-ligand complex structures by specifying pocket residues. In this study, pocket residues were specified as C_*β*_s within 10 Å from the ligand. In addition, the output structure of Umol contains many clashes; therefore, relaxation using OpenMM^26^ is necessary. RoseTTAFold-AA predicts complex structures from the sequence without using ligand-binding position information, similar to AlphaFold3. For Umol and RoseTTAFold-AA, a single inference was performed using default parameters.

## Results

### Evaluation of Predicted Structures

Figure 1 shows the superposition of the HelixFold3 predicted structure and crystal structures, and Figure 2 shows the ligand docking results for Umol, RoseTTAFold-AA, and HelixFold3. The superposition of the HelixFold3 predicted holo structure and the crystal structure generally overlapped. For all targets, the average ligand RMSD was less than 2 Å, indicating that HelixFold3 provides accurate ligand docking. However, for JNK1 and P38, the ligand was slightly shifted, and the centroid distance was larger than that of the other targets. Umol showed ligand RMSDs greater than 2 Å for 4 targets and RoseTTAFold-AA for 5 targets, thus HelixFold3 had the highest ligand docking performance. Umol tends to have smaller centroid distances than RoseTTAFold-AA, likely because Umol specifies pocket residues.

**Figure 1:**
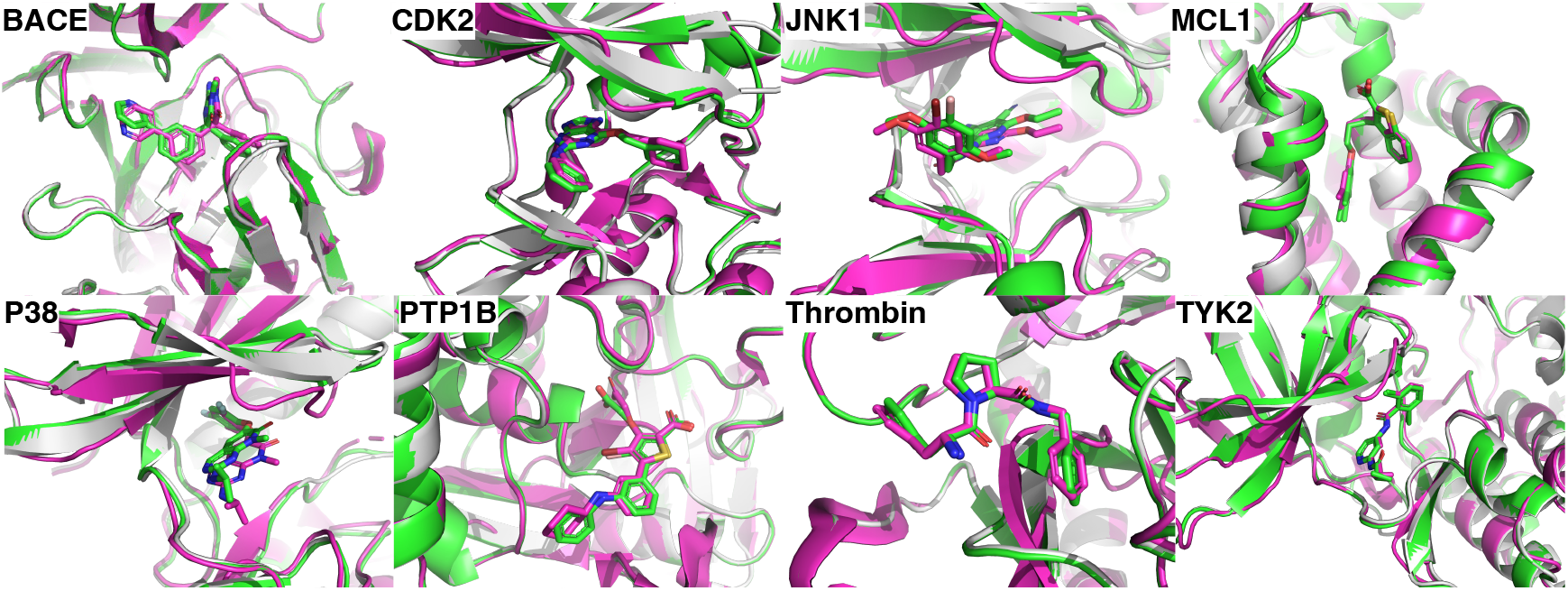
Superposition of the HelixFold3 predicted holo structure (green), apo sructure (white) and crystal structure (magenta).

**Figure 2:**
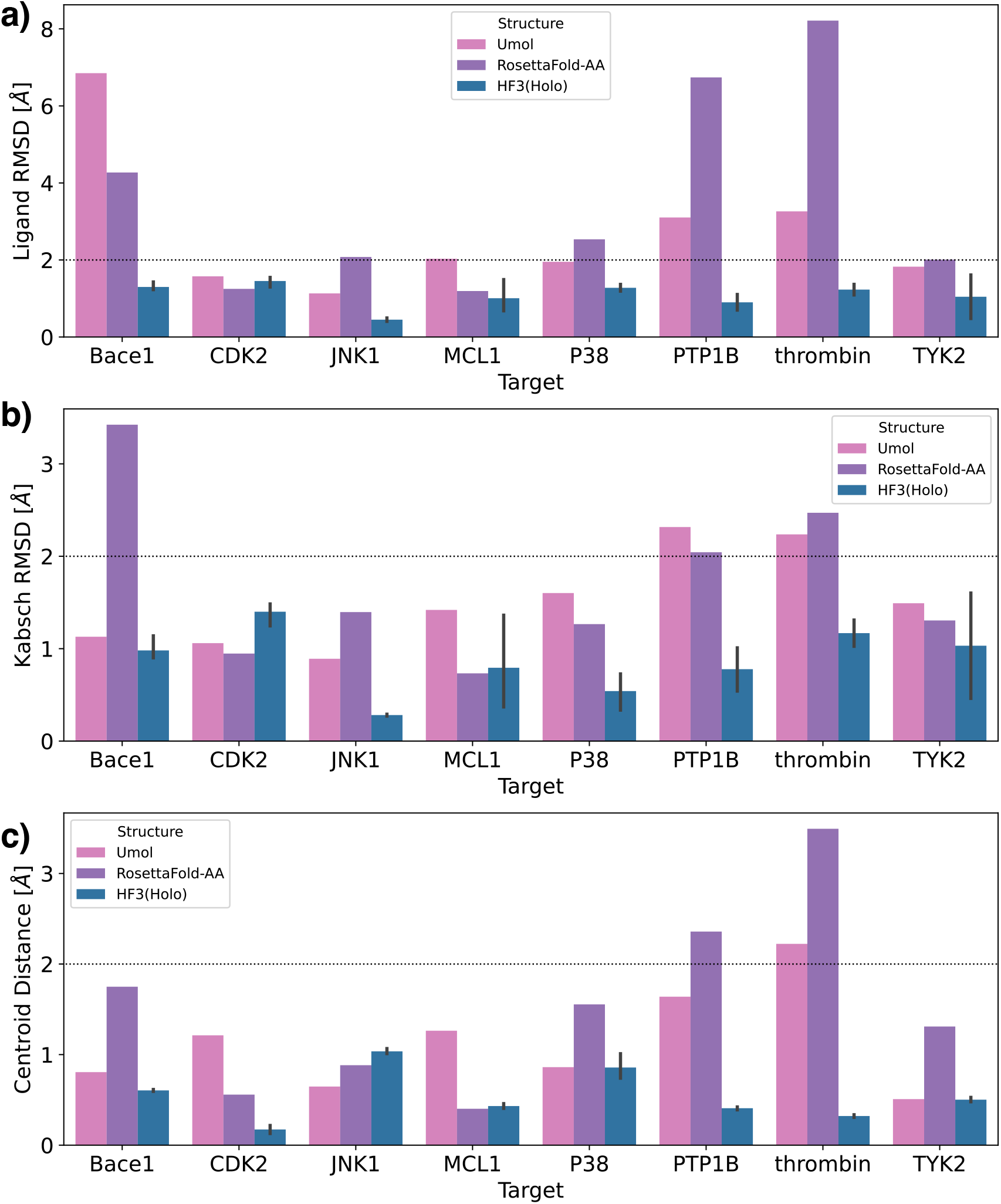
Ligand docking results for protein-ligand complex prediction methods. a) Ligand root mean square deviation (L-RMSD). b) Kabsch RMSD. ^27^ c) Centroid distance. L-RMSD is the root mean square error between the atoms of the predicted ligand and the crystal ligand. Kabsch RMSD is the minimum RMSD when translating and rotating the predicted ligand. Centroid distance is the distance between the centroids of the predicted ligand and the crystal ligand.

Figure 3 shows the global RMSD and binding site RMSD of HelixFold3’s holo and apo structures and existing structure prediction methods. For all targets except JNK1, the global and binding site RMSDs of HelixFold3 predicted structures were the same or lower than those of ColabFold, RoseTTAFold-AA, and Umol, with protein-side structures predicted with high accuracy. HelixFold3, a reproduction of AlphaFold3, not only shows higher ligand docking performance than existing methods but also greatly surpasses conventional methods in binding site prediction accuracy. We compared the prediction accuracies of HelixFold3’s holo and apo structures. In terms of global RMSD, the holo structure had higher accuracy than the apo structure for all targets except BACE and JNK1. In terms of binding site RMSD, the holo structure had significantly higher accuracy than the apo structure for CDK2, MCL1, P38, and PTP1B, and was approximately the same for other targets. This suggests that the holo structure’s prediction performance is comparable to or better than the apo structure, achieving more accurate binding site predictions by utilizing ligand information. However, the binding site RMSDs for JNK1 and P38 were greater than 2 Å for all modeling methods, indicating that although the ligand pose can be predicted, the corresponding pocket structure cannot be accurately predicted. Further improvements in structure prediction methods are expected in these cases.

**Figure 3:**
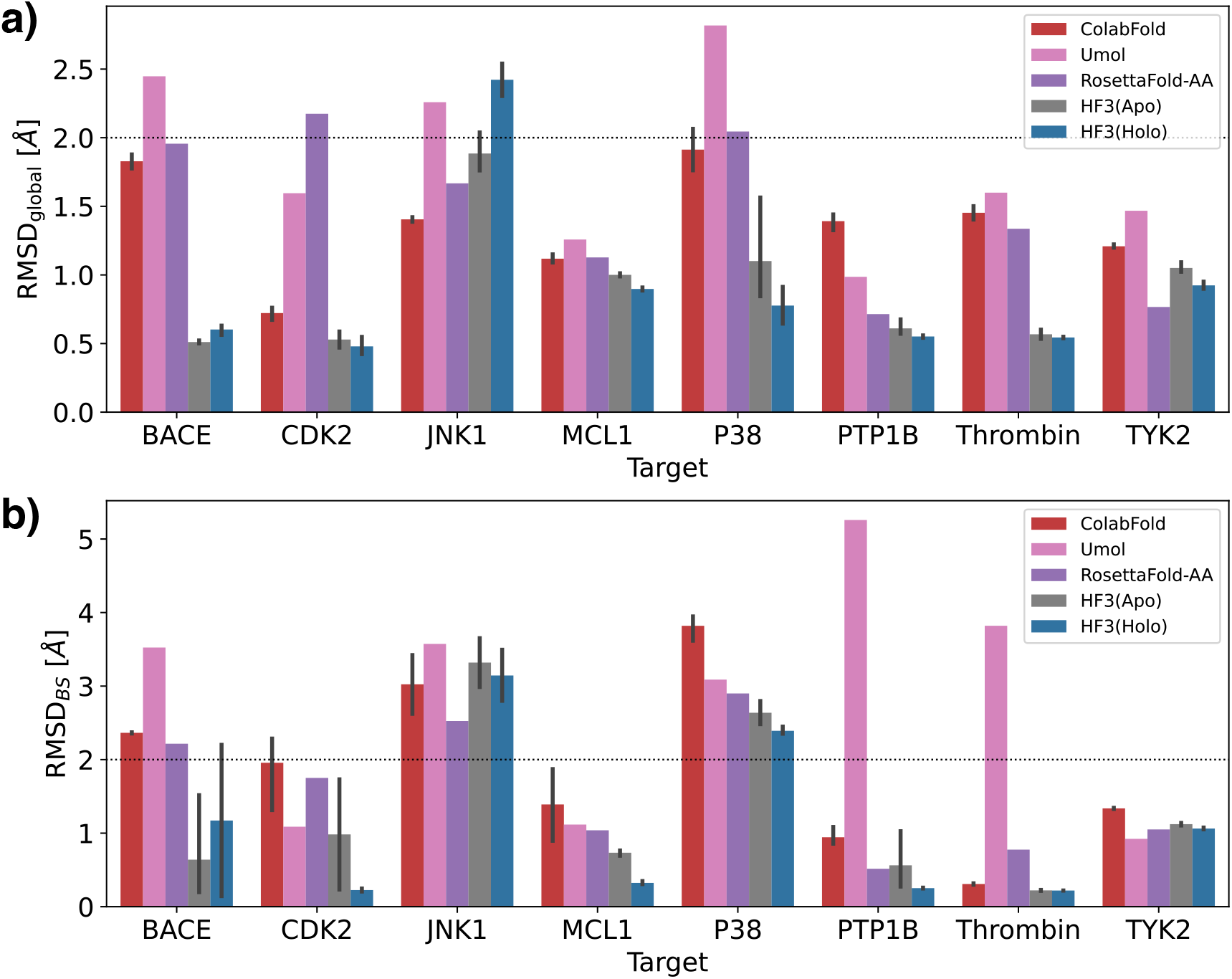
Comparison of predicted structures of ColabFold and HelixFold3’s holo and apo structures with crystal structures. a) Global RMSD. b) binding site RMSD. The RMSD values for the global and binding sites were calculated with only the C*α* atoms. Binding sites are defined as residues within 5 Å of the ligand in the reference structure.

### Evaluation of FEP Performance

Next, we compared the prediction accuracy of binding free energies by Flare FEP using HelixFold3’s predicted holo and apo structures with those using the crystal structure. Figure 4 shows scatter plots of the predicted Δ*G* and experimental Δ*G* for the crystal structure and HelixFold3’s apo and holo structures. Figure 5 shows the mean unsigned error (MUE), Kendall’s *τ*, and Pearson’s correlation coefficient *R*^2^ of each structure. Note that when using the crystal structure of PTP1B, large errors occurred in some perturbation calculations, resulting in significantly low values of Kendall’s *τ* and Pearson’s correlation coefficients *R*^2^. Except for the results of PTP1B using the crystal structure and MCL1’s apo structure, the 90% confidence intervals overlapped, indicating that FEP calculations using HelixFold3 predicted structures can predict binding free energies with the same accuracy as the crystal structure, regardless of apo or holo structure. Even when the pocket structure differs between the crystal and predicted structures, such as JNK1 and P38, the prediction accuracy, including *R*^2^, is the same or higher than that of the crystal structure. For the 4 targets of MCL1, P38, TYK2, and BACE, *R*^2^ tended to be higher for a higher binding-site RMSD. As an exception, for PTP1B, the binding RMSD was smaller for the holo structure, whereas *R*^2^ was higher for the apo structure. This may be because the PTP1B derivative series includes those with large molecular scaffold changes; therefore, the pocket fit to the crystal ligand does not necessarily match the complex structures of the other derivatives. In summary, higher binding site prediction accuracy generally leads to better FEP performance. Therefore, in many cases, holo structure prediction using HelixFold3 is beneficial for FEP calculations.

**Figure 4:**
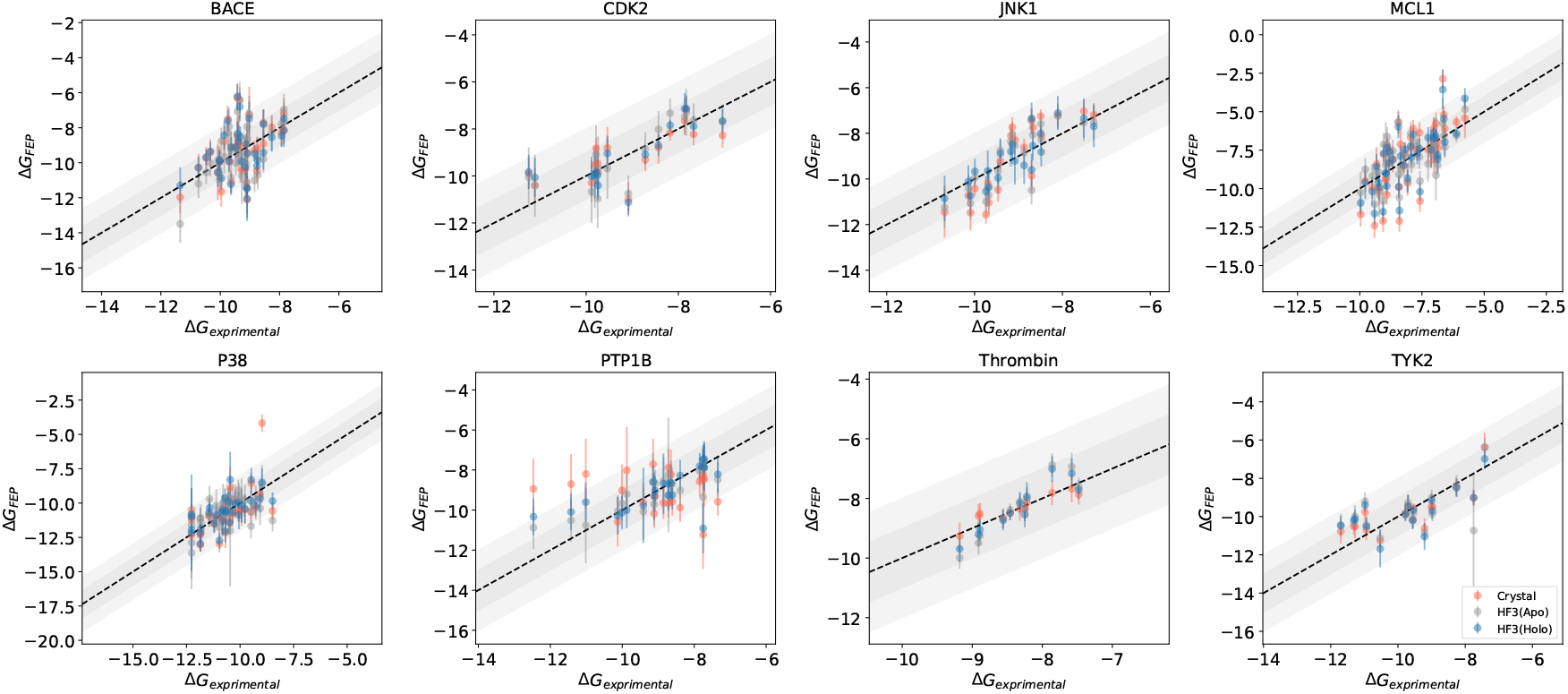
Scatter plots of experimental Δ*G* and predicted Δ*G* for the apo and holo structures of HelixFold3 and the crystal structure. The dark gray area indicates a range of *±* 1.0 kcal/mol, and the light gray area indicates a range of *±* 2.0 kcal/mol.

**Figure 5:**
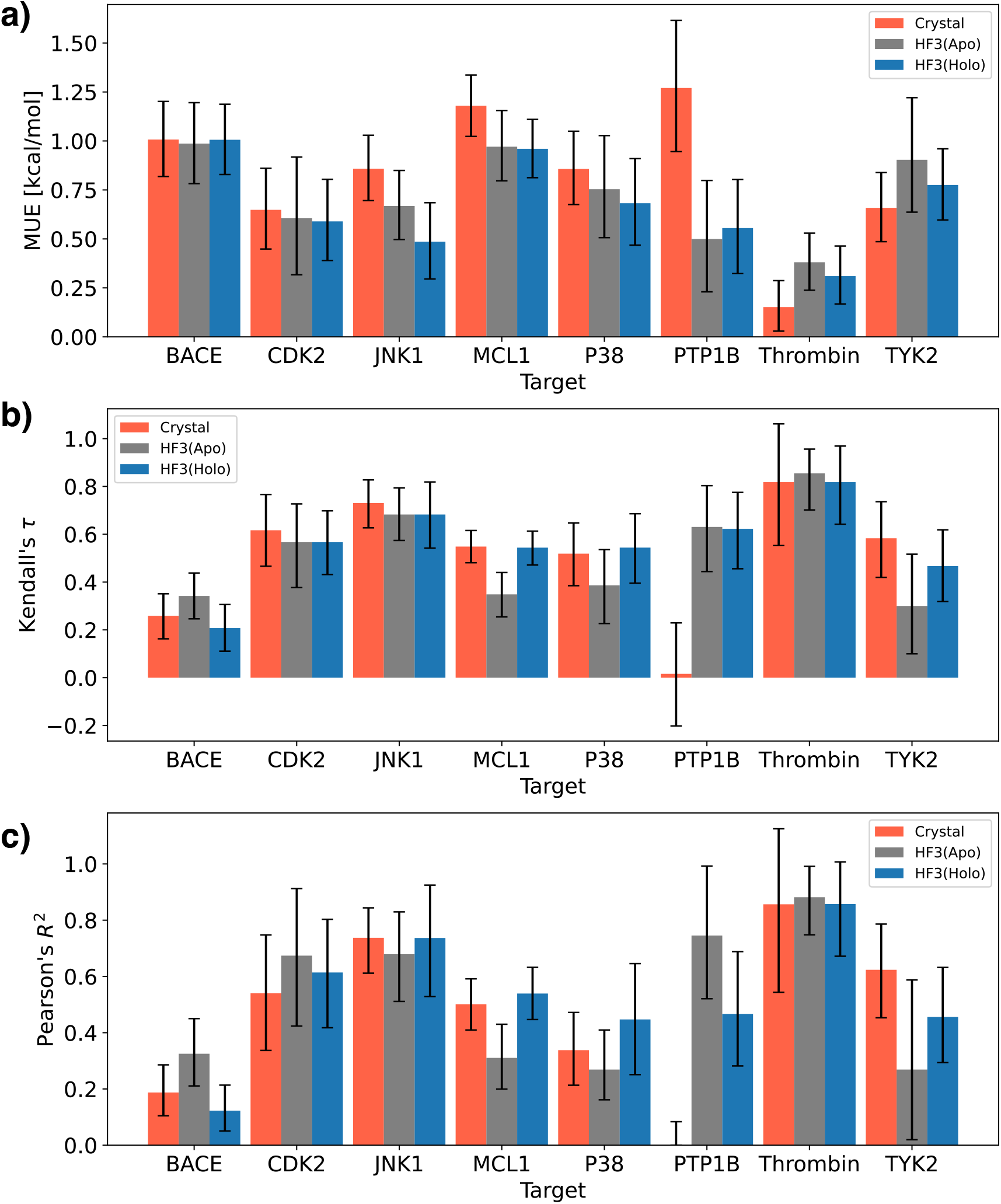
Performance of Flare FEP for the apo and holo structures of HelixFold3 and the crystal structure. a) Mean unsigned error (MUE). b) Kendall’s *τ* . c) Pearson’s correlation coefficient *R*^2^. Error bars indicate 90% confidence intervals calculated by 10,000 bootstrap samplings.

### Evaluation of Predicted Structures in Derivative Sets

Figure 6 shows the distribution of ligand RMSDs of HelixFold3’s predicted structures for the derivative set. The figure confirms that for JNK1 derivatives, there are cases where the ligand RMSD exceeds 2 Å (8 out of 21 JNK1 derivatives). Figure 7 shows the superposition of the complex structure predicted by HelixFold3 with the lowest ligand RMSD in the JNK1 derivative set and the crystal structure. As shown in Figure 1, the binding position of the derivative ligand was generally correct; however, the binding pose was inaccurate. For other targets, there were no derivatives with ligand RMSD significantly exceeding 2 Å, indicating that HelixFold3 can generally perform accurate ligand docking, even for ligands not included in the training data.

**Figure 6:**
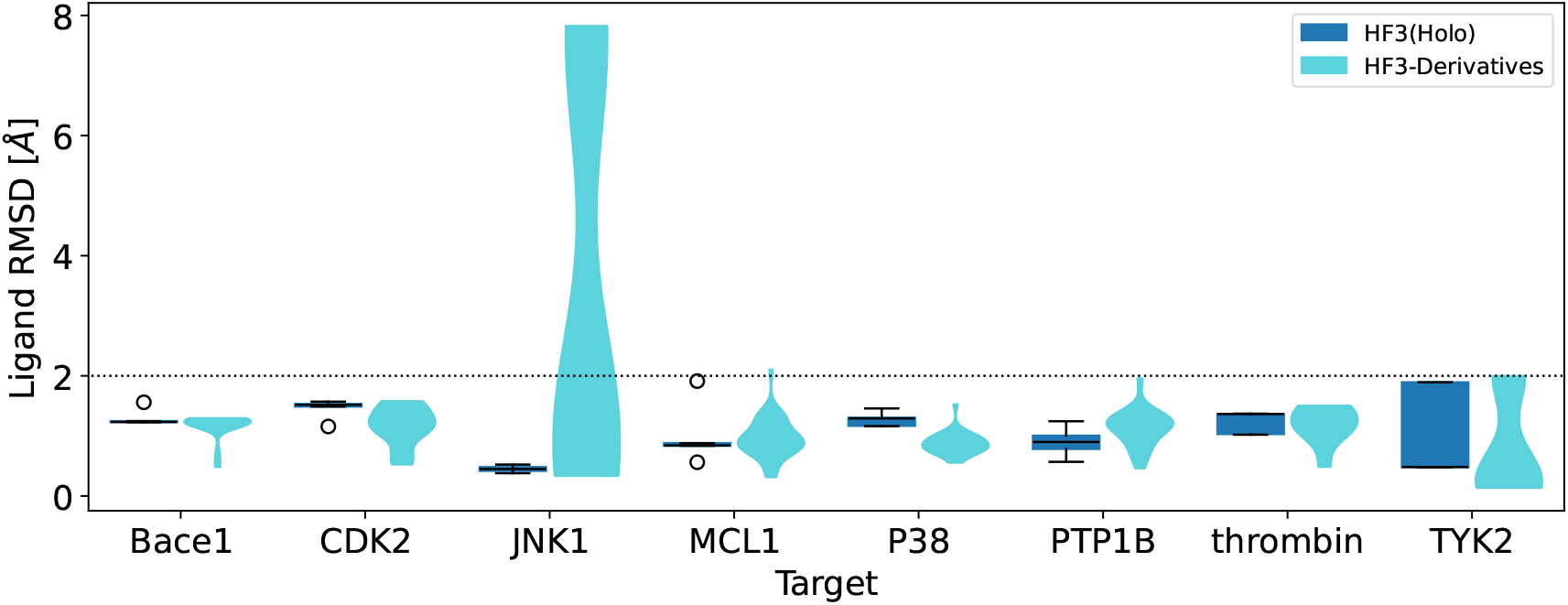
Distribution of ligand RMSD of predicted structures for the derivative set. HF3(Holo) shows the distribution of 5 predicted structures with the crystal ligand as a box plot, and HF3-Derivatives shows the distribution for the entire derivative set of Wang *et al*.’s benchmark set as a violin plot. Ligand RMSD is calculated for the maximum common substructure between the crystal ligand and the derivative.

**Figure 7:**
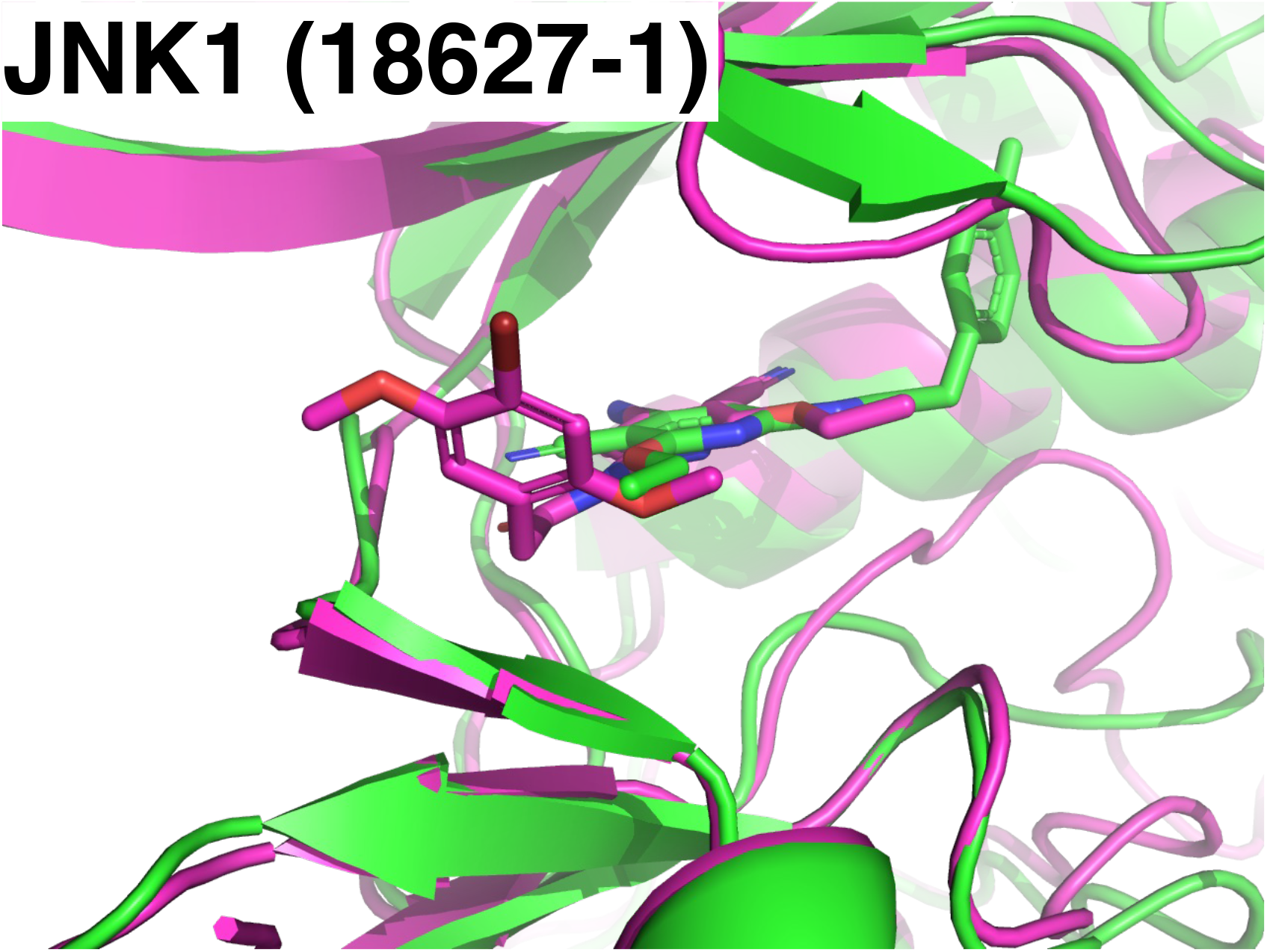
Superposition of the predicted complex structure by HelixFold3 with the worst ligand RMSD in the JNK1 derivative set (green) and the crystal structure (purple). The ligand is 18627-1.

Figure 8 shows the relationship between ligand RMSD and experimental binding free energy for JNK1. The figure shows that the binding free energy of the group that predicted the correct ligand position tended to be lower than that of the group that did not. This suggests the ligand pose accuracy may contribute to the prediction accuracy of binding free energy; however, in this dataset, incorrect ligand poses were not predicted, except for the JNK1 derivative set. Therefore, further verification is needed to determine if a general statement can be made regarding the relationship between ligand pose error and binding free energy.

**Figure 8:**
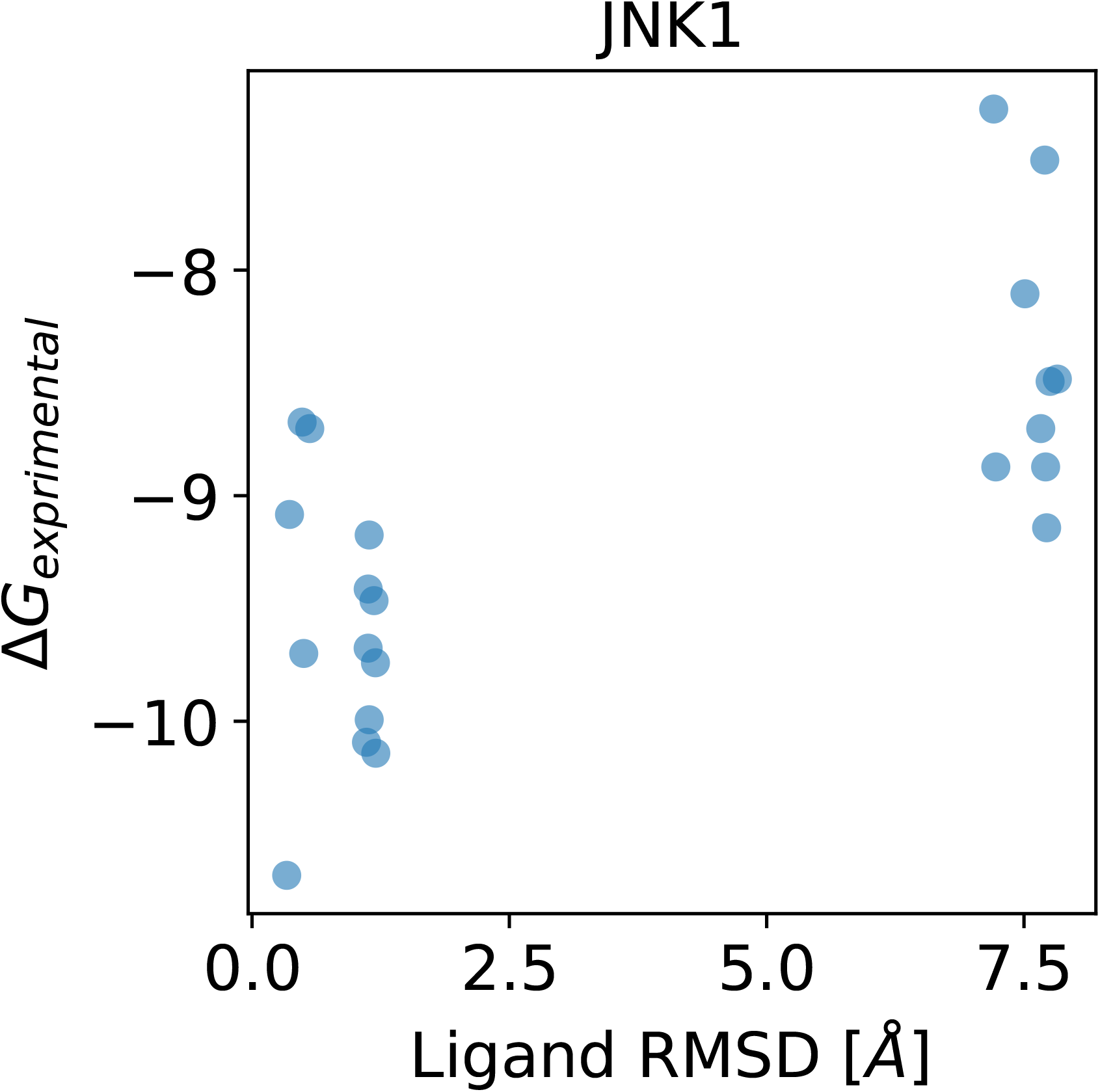
Relationship between ligand RMSD and experimental binding free energy for the JNK1 derivative set.

Table 2 shows the correlation between HelixFold3’s ranking score and the experimental binding free energy. The table confirms that for some targets, there is a certain correlation between the ranking score and binding free energy. This suggests that, as Lu *et al*.^28^ indicated, the ranking score can predict the impact of mutations on protein-protein interactions and may also predict binding free energy for derivative sets in some cases.

**Table 2:** Performance of HelixFold3’s ranking score for derivative sets.

| Target | Pearson’s $R^2$ | Kendall’s $\tau$ |
| --- | --- | --- |
| BACE | 0.24 | 0.33 |
| CDK2 | 0.33 | 0.50 |
| JNK1 | 0.38 | 0.44 |
| MCL1 | 0.08 | 0.24 |
| P38 | 0.00 | -0.06 |
| PTP1B | 0.00 | -0.02 |
| Thrombin | 0.63 | 0.78 |
| TYK2 | 0.17 | 0.22 |

Figure 9 shows the global RMSD and binding-site RMSD of HelixFold3’s predicted structures superimposed on crystal structures for the derivative set. The figure confirms that for JNK1, there are complexes with binding-site RMSDs of less than 2 Å. While ligand RMSDs tend to worsen, binding-site RMSDs improve, resulting in a reversal phenomenon. The diversity in the predicted structures may result from slight changes in the derivatives’ structures from the crystal ligand, resulting in predictions close to the crystal structure. For other targets, the binding site prediction accuracy did not change significantly when a crystal ligand was used. Therefore, HelixFold3 has a certain robustness in predicting binding sites, even for derivatives not included in the training data.

**Figure 9:**
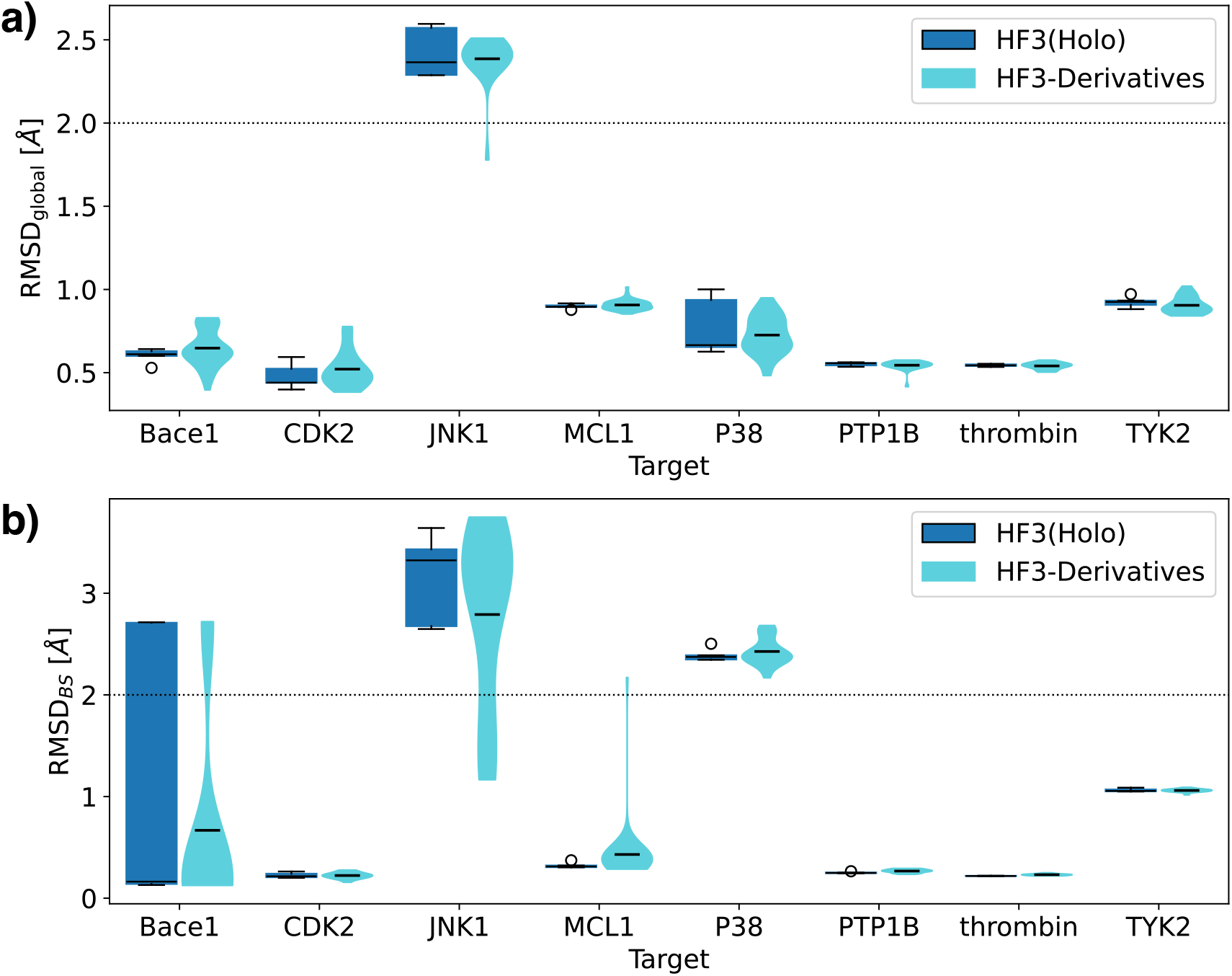
Comparison of predicted structures and crystal structures for the derivative set. Global RMSD. b) binding site RMSD. HF3(Holo) shows the distribution of 5 predicted structures with the crystal ligand as a box plot, and HF3-Derivatives shows the distribution for the entire derivative set of Wang *et al*.’s benchmark set as a violin plot.

### Evaluation of FEP Performance on Derivative Sets

Finally, for these predicted structures, a single structure with a different conformation from the crystal ligand’s predicted structure was selected to perform FEP calculations and evaluate the binding free energy prediction accuracy. First, to select a structure different from the crystal ligand’s predicted structure, the distance matrix between the pocket residues was compressed into 2 dimensions using Multidimensional Scaling (MDS) and classified into 3 clusters using hierarchical clustering. Because of computational resource limitations, from the 2 clusters that did not contain the predicted structure with the crystal ligand, only one structure closest to the cluster center of the larger cluster was selected for FEP. Table 3 lists the selected derivative for each target, its ligand RMSD, and the binding site RMSD. The receptors of these predicted complex structures were extracted and superimposed onto the crystal structure for FEP calculations.

**Table 3:** Selected derivatives and their ligand RMSDs and binding site RMSDs.

| Target | Ligand | Ligand RMSD [Å] | RMSD<sub>BS</sub> [Å] |
| --- | --- | --- | --- |
| BACE | CAT-17c | 1.26 | 0.22 |
| CDK2 | 31 | 1.11 | 0.22 |
| JNK1 | 18638-1 | 1.11 | 1.17 |
| MCL1 | 39 | 0.75 | 0.53 |
| P38 | p38a_2g | 0.68 | 2.65 |
| PTP1B | 23484 | 1.11 | 0.25 |
| Thrombin | 1b | 1.47 | 0.23 |
| TYK2 | jmc_30 | 0.20 | 1.02 |

Figure 10 shows the performance of Flare FEP for the predicted structures of the crystal ligand and selected derivative. The figure shows that when using the MCL1 derivative, the MUE was significantly worse than when using the crystal ligand, and there was no significant difference between Kendall’s *τ* and Pearson’s *R*^2^.

**Figure 10:**
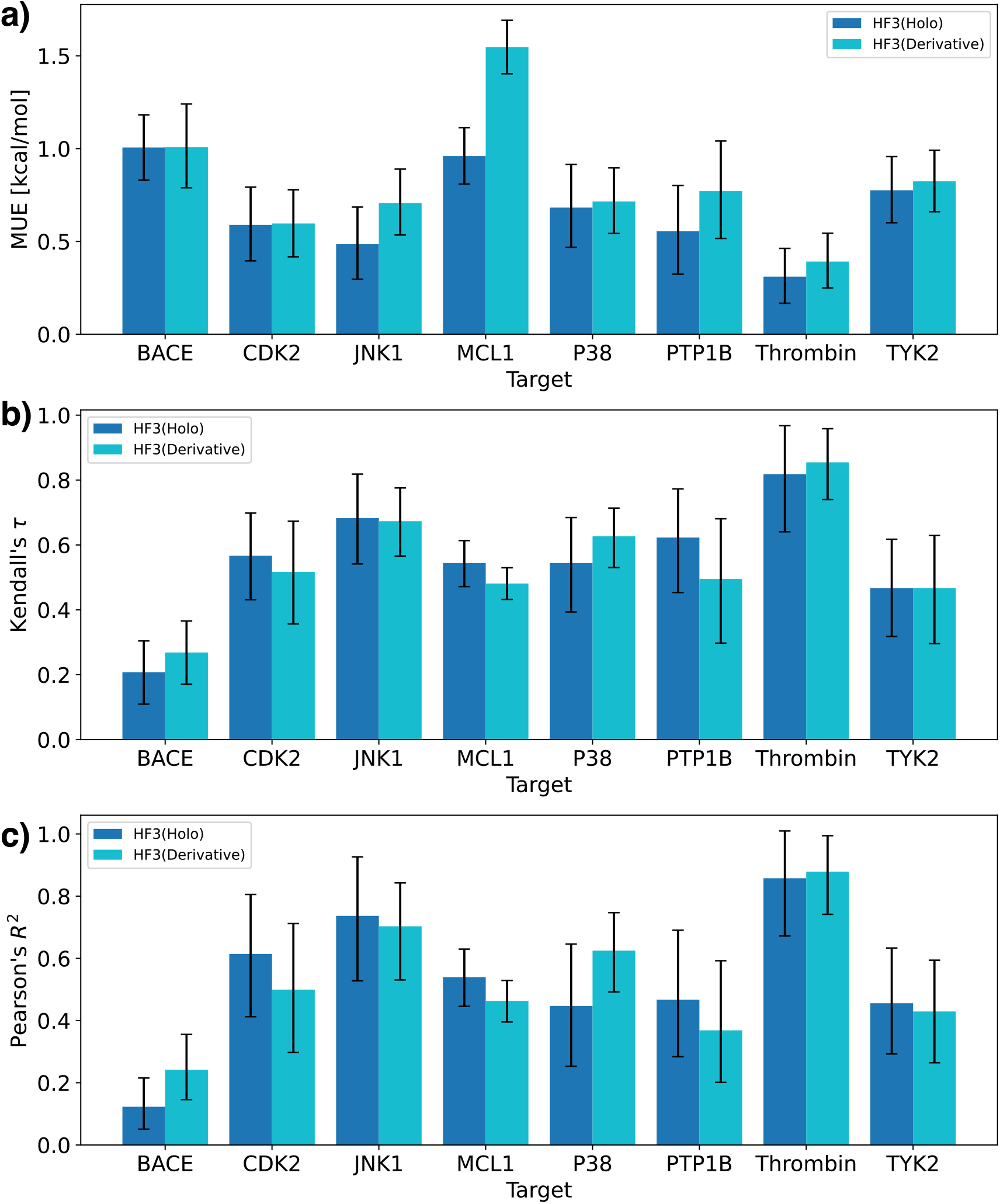
Performance of Flare FEP for the predicted structures selected from the derivative set (HF3(Holo) is the result for the crystal ligand, and HF3(Derivative) is the result for the predicted structure of the selected derivative). a) Mean unsigned error (MUE). b) Kendall’s *τ* . c) Pearson’s correlation coefficient *R*^2^.

In addition, for other targets, although there were overlaps in the confidence intervals, in many cases the MUE was smaller when using the crystal ligand, suggesting a tendency toward higher prediction accuracy with the crystal ligand. In conclusion, while using the predicted crystal ligand complex structure leads to higher prediction accuracy, sufficiently accurate FEP calculations can still be performed using derivative ligand complex structures. An important limitation of this study is that the target structures of the Wang *et al*. dataset are known, and whether HelixFold3 can predict accurate pocket structures for targets without experimentally solved structures or FEP performance in such cases has not been evaluated. In addition, in this study, the ligand poses for FEP calculations were those of the crystal; however, in cases where experimental complex structures are not available, FEP calculations need to be performed from the predicted ligand poses, and the impact on FEP calculations using the predicted poses also needs to be evaluated in the future.

## Conclusion

In this study, using the state-of-the-art structure prediction model HelixFold3, we compared the predicted holo and apo structures of protein-ligand complexes and evaluated the prediction accuracy of binding free energies by Flare FEP using these predicted structures. HelixFold3 can predict pocket structures more accurately than ColabFold and existing proteinligand complex prediction methods. In many cases, the holo structure can model pocket structures more accurately than the apo structure. In the performance evaluation by Flare FEP, it was found that using HelixFold3 predicted structures, binding free energies could be predicted with accuracy comparable to that using crystal structures. There was a tendency for better prediction performance in FEP with smaller pocket RMSDs. Furthermore, validation of the derivative set suggested the robustness of HelixFold3, as it can generally predict complex structures with ligands not included in the training data, and FEP calculations using complex structures with derivatives can also predict binding free energies with sufficient accuracy. It is also suggested that the ranking score and ligand RMSD could be useful for estimating binding free energies. Future studies should assess the accuracy of HelixFold3’s modeled structures in compound docking and explore methods for modeling appropriate pocket structures using HelixFold3 when experimental structural information is limited. The demonstrated utility of HelixFold3’s complex structures for FEP calculations indicates that these predicted structures could expedite early stages of small molecule drug discovery, such as lead optimization.

## Data Availability

### Supporting Information Available

The information of each predicted structure and results is available on GitHub, https://github.com/ohuelab/helixfold3-benchmark-fep. The Flare project files (.flr) are available on Google Drive, https://drive.google.com/file/d/1Mckue-rnBxlV06fZrsnNMcXiY0rS0kKm.

## Author Information

### Corresponding Author

Masahito Ohue - School of Computing, Institute of Science Tokyo, Yokohama, Kanagawa 226-8501, Japan.

